# A predictive coarse-grained model for position-specific effects of post-translational modifications on disordered protein phase separation

**DOI:** 10.1101/2020.06.12.148650

**Authors:** T. M. Perdikari, N. Jovic, G. L. Dignon, Y. C. Kim, N. L. Fawzi, J. Mittal

## Abstract

Biomolecules undergo liquid-liquid phase separation (LLPS) resulting in the formation of multicomponent protein-RNA membraneless organelles in cells. However, the physiological and pathological role of post translational modifications (PTMs) on the biophysics of phase behavior is only beginning to be probed. To study the effect of PTMs on LLPS *in silico*, we extend our transferable coarse-grained model of intrinsically disordered proteins to include phosphorylated and acetylated amino acids. Using the parameters for modified amino acids available for fixed charge atomistic forcefields, we parameterize the size and atomistic hydropathy of the coarse-grained modified amino acid beads, and hence the interactions between the modified and natural amino acids. We then elucidate how the number and position of phosphorylated and acetylated residues alter the protein’s single chain compactness and its propensity to phase separate. We show that both the number and the position of phosphorylated threonines/serines or acetylated lysines can serve as a molecular on/off switch for phase separation in the well-studied disordered regions of FUS and DDX3X, respectively. We also compare modified residues to their commonly used PTM mimics for their impact on chain properties. Importantly, we show that the model can predict and capture experimentally measured differences in the phase behavior for position-specific modifications, showing that the position of modifications can dictate phase separation. In sum, this model will be useful for studying LLPS of post-translationally modified intrinsically disordered proteins and predicting how modifications control phase behavior with position-specific resolution.

**Statement of Significance:** Post-translational modifications are important regulators of liquid-liquid phase separation (LLPS) which drives the formation of biomolecular condensates. Theoretical methods can be used to characterize the biophysical properties of intrinsically disordered proteins (IDPs). Our recent framework for molecular simulations using a Cα-centered coarse-grained model can predict the effect of various perturbations such as mutations (Dignon et al. *PloS Comput. Biol*, 2018) and temperature (Dignon et al, *ACS Cent. Sci.*, 2019) on LLPS. Here, we expand this framework to incorporate modified residues like phosphothreonine, phosphoserine and acetylysine. This model will prove useful for simulating the phase separation of post-translationally modified IDPs and predicting how position-specific modifications can control phase behavior across the large family of proteins known to be phosphorylated and acetylated.

## Introduction

Many cellular processes depend on the formation of membraneless nuclear and cytoplasmic assemblies known as membraneless organelles or biomolecular condensates(1–3). Lacking a phospholipid membrane, these biomolecular condensates(4) can respond rapidly to environmental changes, forming cellular compartments concentrating specific protein(5) and nucleic acids(6). The formation of many membraneless organelles appears to be driven by interactions between proteins containing intrinsically disordered regions (IDRs)(7). The detailed molecular interactions mediating these membraneless organelles may include cation-π(8), sp^2^/π(9), hydrogen bonding, salt bridges and hydrophobic interactions(10–12). Importantly, recent efforts have demonstrated that IDR LLPS and, hence, the formation of membranelles organelles can be regulated by a plethora of factors such as salt concentration, pH, RNA(13, 14), ATP(15), temperature(16) and post translational modifications (PTMs)(17, 18).

PTMs enrich the repertoire of the 20 natural amino-acids and have been shown to be an important potential means for modulating phase-separation(18). Two of the most frequently occurring eukaryotic PTMs(19) are phosphorylation(20) and acetylation(21) which covalently attach a phosphoryl group or an acetyl group, respectively, to select amino acids(22). These modifications can reduce the binding affinity of RNA(23), impair enzyme activity(24), decrease aggregation propensity of fibril-forming segments(25) or trigger the self-assembly of RNA granules(26). Importantly, phosphorylation and acetylation both change the charge of the associated amino acid. Phosphorylated serine and threonine have a net −2 charge at neutral pH compared to no net charge for unmodified serine and threonine. Conversely, acetylation neutralizes the +1 charge that lysine has at physiological pH. Because IDRs generally contain many PTM sites which may be simultaneously modified(27), this can dramatically alter the net charge and charge patterning in ways that impact chain properties, phase separation, and aggregation. Hyperphosphorylation and hyperacetylation can lead to large changes in protein charge distribution, and hence conformation, and are associated with aggregation of amyloidogenic proteins in neurodegenerative diseases(21, 28, 29).

Although *in vitro* studies have demonstrated the effect of different PTMs on LLPS, studying the effect of each possible combination of PTMs is laborious and difficult. Furthermore, it is challenging to have site-specific control to reproduce modification patterns that are deposited on IDRs *in vivo*. Therefore, computational studies can be an effective approach to screen a large number of PTM patterns and predict their role on regulating the molecular properties and phase separation of proteins(30, 31). All-atom explicit solvent simulations readily incorporate post-translationally modified amino acids(32–36) and have been used to probe the contacts leading to phase separation(37, 38). However, the ability of atomistic force-fields to accurately capture the properties of IDRs depends strongly on parameterization(39, 40). Additionally, simulations of sufficiently long length and time scales to predict phase separation behavior (e.g. saturation concentration) are beyond current computational capability for fully atomistic simulations with explicit water. In contrast, coarse-grained (CG) models(41), in particular those in which peptides are represented as chains of single beads and solvation effects are represented implicitly, have been used to efficiently investigate the phase-behavior of large macromolecules(42, 43). Most of current CG modeling approaches capable of studying IDRs are currently parameterized only for the 20 natural amino acids(44) while others are suited specifically to probe phosphoregulation of folded domains(45, 46). Therefore, there is a need for transferable approach to elucidate the effect of PTMs on the biophysical properties and phase-behavior of IDRs associated with the assembly of biomolecular condensates.

Here we expand our previous amino acid resolution coarse-grained model(44) to incorporate PTMs compatible with the procedure used to obtain the parameters for the 20 natural unmodified amino acids. To investigate how the position and the number of modified sites can alter the phase behavior of IDRs, we perform *in silico* studies of two IDRs: the N-terminal low-complexity (LC) domain of RNA-binding protein <u>Fu</u>sed in Sarcoma (FUS LC) which is rich in serine, tyrosine, glutamine, and glycine residues and contains 12 known phosphorylation sites modified by DNA-dependent protein kinase(47) and the first disordered region (IDR1) of the DEAD box RNA helicase 3, X-linked (DDX3X) with 10 lysine residues known to be acetylated(48). We test how the post-translational state of each molecule modulates IDR compactness and LLPS in a sequence-specific manner. Using coarse-grained molecular simulations, we examine the effect of increasing number of modified residues and test the hypothesis that position-specific patterning of PTMs can alter the collapse and phase behavior of IDR sequences. Finally, we test the ability of the model to reveal the origin of experimentally known position-specific effects on phase separation.

## Results

### Expanding the HPS model to incorporate phosphorylation and acetylation

To extend our primary sequence-specific coarse-grained model to study PTMs, we employed the original philosophy of computing coarse-grained interaction parameters from the size, net charge, and atomic hydropathy of the post translationally modified amino acids. Our previous coarse-grained representation of the protein represents every natural amino acid by a single bead of unique mass, charge, size and hydropathy parameter that were computed from a fully-atomistic fixed charge forcefield parameters for each residue type(44, 49, 50). Using previously published AMBER force-field parameters for modified amino acids(34), we computed coarse-grained parameters for serine, threonine, and tyrosine phosphorylation and lysine acetylation (**Table S1**). In principle, these calculations can be applied to any library of non-canonical amino-acids with known structure/charge parameters to aid the high-throughput screening of proteins with unknown PTM-regulated phase separation.

To test the effects of these PTMs on coarse grained models of IDRs, we used the HPS-PTM model to characterize the properties of two modified intrinsically disordered regions: phosphorylated FUS LC (FUS LC^12pS/pT^) and acetylated human DDX3X IDR1 protein (DDX3X IDR1^10acK^). Single-chain coarse-grained simulations were conducted for 1 μs at a range of temperatures (150-600 K) using replica exchange molecular dynamics (REMD) to enhance sampling(51). Unmodified FUS LC is nearly uncharged (containing only two charged residues) but can be phosphorylated by DNA-dependent protein kinase at 12 unique sites, displaying either SQ or TQ motif(47). Here, we begin by testing the effect of hyperphosphorylation at all 12 sites (FUS LC^12pS/pT^). In the hyperphosphorylated state, FUS LC^12pS/pT^ is more extended than the unmodified state, having a much larger radius of gyration (*R*_g_) at the reference temperature (300K) (**Figure 1A**). Hyperphosphorylation also reduces *T_θ_* (**Figure S1A,C)**, the temperature at which attractive intramolecular interactions are balanced by repulsive excluded volume interactions(52). Both of these reflect experimental observations that phosphomimetic mutants of FUS are more extended in solution and are less prone to phase separate(25).

**Figure 1:**
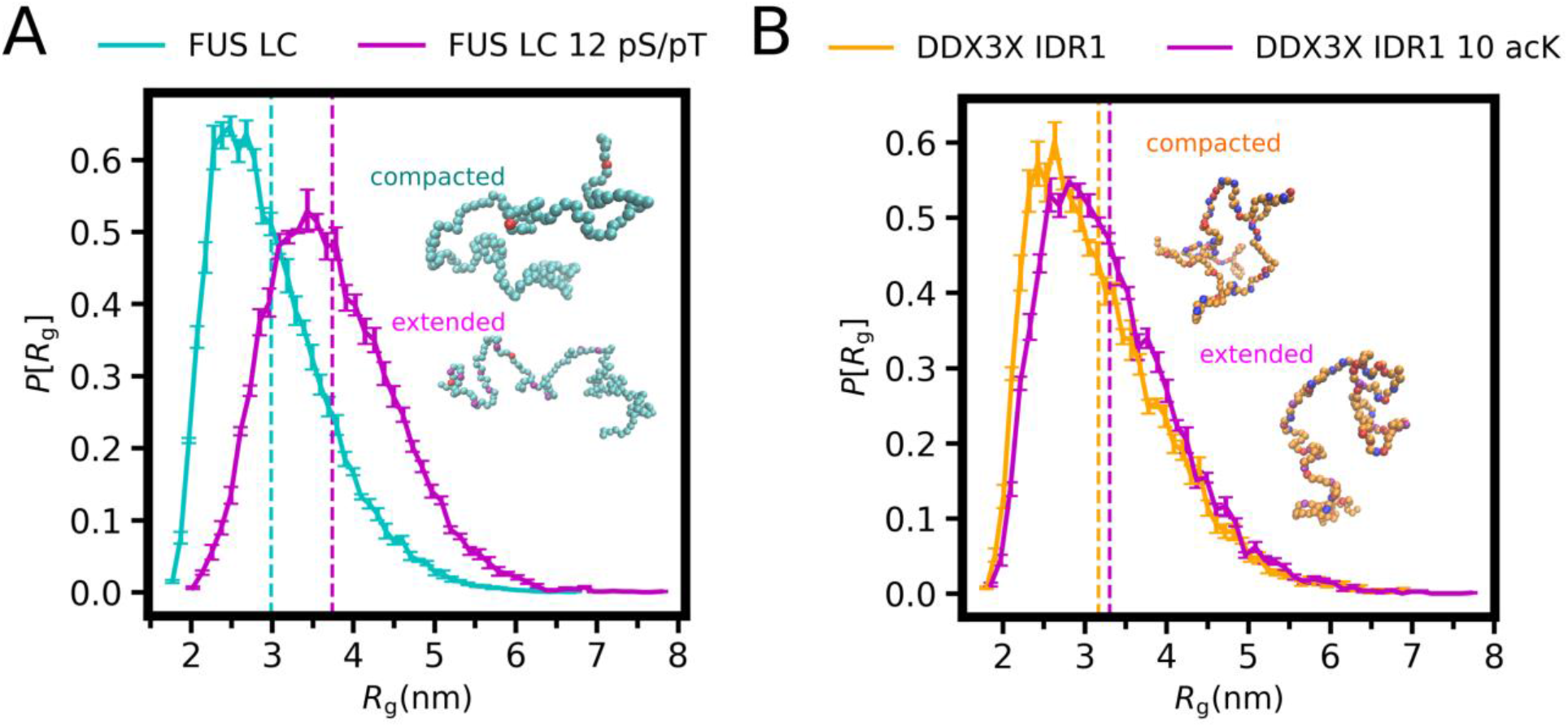
Expanding the HPS model to incorporate phosphorylation and acetylation. **A)** Distribution of ensemble radius of gyration of FUS LC (cyan) and FUS LC 12 pS/pT (purple). **B)** Distribution of ensemble radius of gyration of DDX3X IDR1 (orange) and DDX3X IDR1 10 acK (purple). In representative snapshots, the unmodified beads are shown in cyan (FUS LC) or orange (DDX3X IDR1), the negatively and positively charged in red and blue, respectively, and the modified sites in purple.

Next, we tested the effect of acetylation using single-chain REMD simulations of DDX3X IDR1 and DDX3X IDR1^10acK^. At the reference 300K temperature, we found that the modified state is also more expanded than the unmodified state, but the difference is considerably less than that of FUS LC^12pS/pT^ from unmodified FUS LC (**Figure 1B**). Similarly, *T_θ_* is only slightly decreased (**Figure S1B, D**), consistent with the small change in *R*_g_. This is also consistent with experimental results showing that acetylmimetic DDX3X is less prone to phase separate(48).

We suspect that the large changes in single chain properties for FUS LC hyperphosphorylation compared to DDX3X IDR1 hyperacetylation may be due to the nearly uncharged nature of unmodified FUS LC as opposed to the highly charged polyampholytic character of DDX3X IDR1, where other positively and negatively charged residues are extensively distributed throughout the sequence. Therefore, acetylation of lysine residues across the entire DDX3X IDR does not dramatically alter the intramolecular interactions, which we probe further in the following sections. Taken together, these data demonstrate that we can model changes in biophysical properties of IDRs using this coarse-grained approach.

### Effect of increasing number of post-translational modifications on chain properties of FUS LC and DDX3X IDR1

We next wanted to test the effect of increasing the number of modified sites on the expansion or collapse of the IDRs. The number of possible modification combinations is very large, given by the following equation:

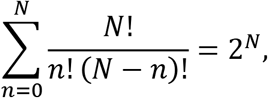

where N is the total number of possible PTM sites, and n is the number of sites that are modified. While there is only one possible state with 0, or N modifications, there are 924 possible states of FUS with 6 modified residues, and 252 possible states of DDX3 with 5 modified residues. We selected a subset of 10 random modification patterns for each number of modifications sites (1≤n≤11 for FUS LC and 1≤n≤9 for DDX3X IDR), and performed simulations on these sequences. We then calculated the sequence charge decoration (SCD) of all sequences to determine the extreme cases where charge patterning was the most, and the least, and ensured that both extremes were included in our list of sequences in order to maximize the expected variance in behavior. SCD is calculated as(53):

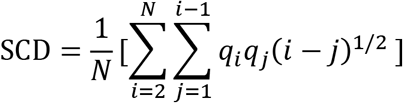

For FUS LC, *R*_g_ at 300 K increases with increasing number of phosphorylation sites (**Figure 2A, B**), going from slightly below 3.0 nm for unmodified to 3.8 nm for FUS LC^12pS/pT^. This large expansion is consistent with increasing polyelectrolytic character due to phosphorylation, adding increasing numbers of negatively charged sites to FUS LC which is otherwise nearly uncharged. It is important to note that a large change in *R*_g_ can be obtained within a fixed number of phosphorylation sites by changing the sequence (due to the variance in *R*_g_ for the different phosphorylation patterns) or by slightly changing the number of phosphorylation sites, which can be attributed to differences in charge patterning, and perhaps also hydropathy patterning (54). We also note that in many cases, the sequence of the lowest possible SCD value for a given number of phosphorylated residues frequently has the one of the lowest *R*_g_ values. However, in the case of the highest possible SCD value, the sequences do not have the highest *R*_g_ value (**Figure 2B**). This could be due to the changes in SCD being very small within these sequences, and the occasional disagreement between SCD and chain dimensions as observed previously (55).

**Figure 2:**
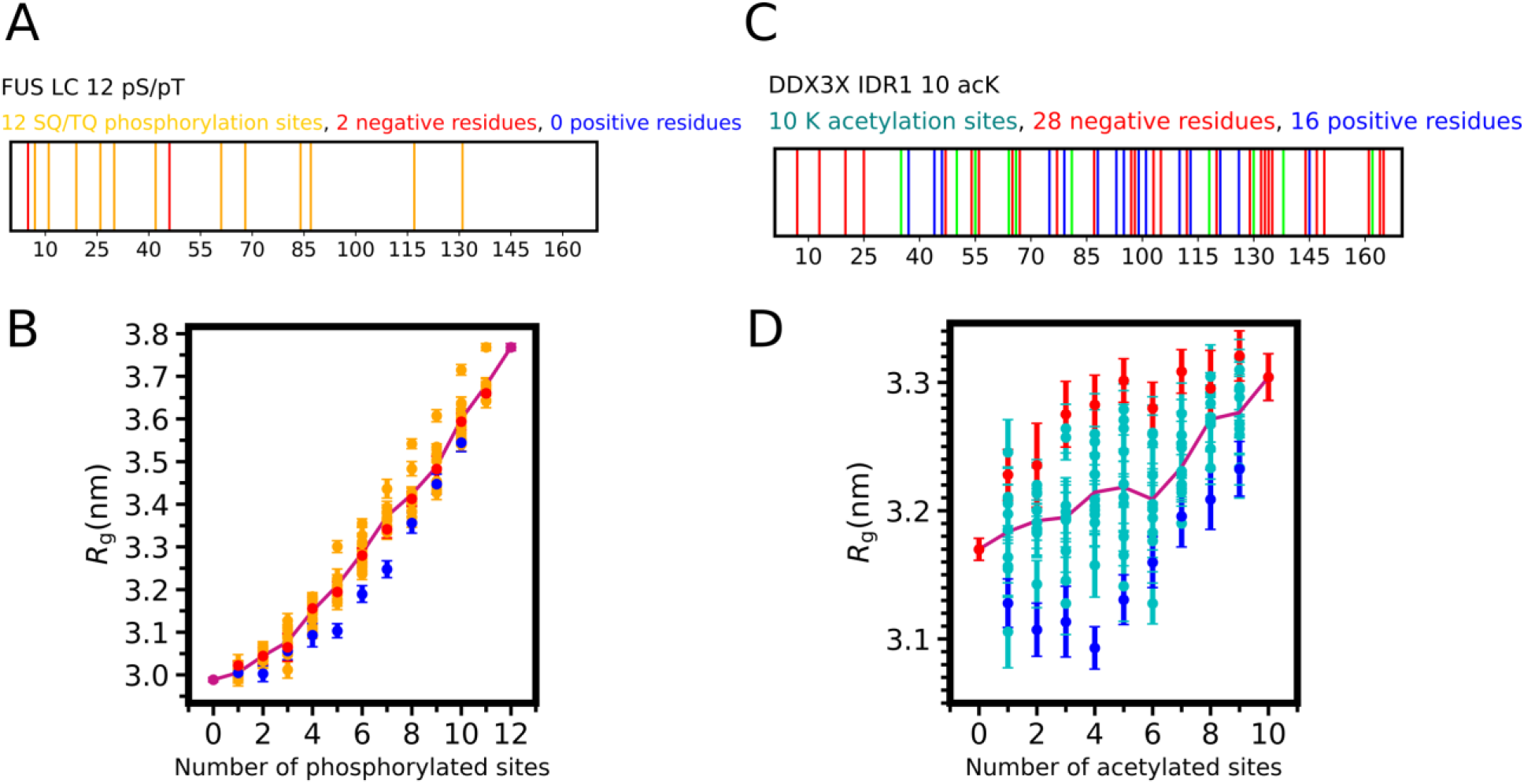
Effect of increasing number of post-translation modifications on chain properties of FUS LC and DDX3X IDR1. **A)** Domain representation of FUS LC 12 pS/pT. Negative residues D5 and D46 are highlighted in red. Phosphorylation sites at 7, 11, 19, 68 (phosphothreonine) and 26, 30, 42, 61, 84, 87, 117, 131 (phosphoserine) are highlighted in orange. **B)** Random phosphorylation patterns expand FUS LC monotonically. Orange data points show the ensemble average *R*_g_ and SEM from 10 equal divisions of the equilibrated ensembles for each variant of 10 randomly selected phosphorylation patterns for a given number of phosphorylation sites in FUS LC. Violet line represents the mean *R*_g_ value from 10 simulations. Red and blue data points show the *R*_g_ of SCD maximum and SCD minimum sequences respectively. **C)** Domain representation of DDX3X IDR1. Negative residues are shown in red, positive residues (other than lysine, i.e. arginine) in blue and lysines in green. **D)** Random acetylation patterns in DDX3X yield to a moderate expansion. Blue data points show the ensemble average *R*_g_ and SEM from 10 equal divisions of the equilibrated ensembles for each variant of 10 randomly selected acetylation patterns for a given number of acetylation sites in DDX3X IDR1. Violet line represents the mean *R*_g_ value from 10 simulations. Red and blue data points show the *R*_g_ of SCD maximum and SCD minimum sequences respectively.

We then probed the chain properties of the DDX3X IDR as a function of lysine acetylation. To quantify the difference in chain properties as a function of increasing number of acetylation modifications, we performed single-chain simulations selecting 10 random acetylation patterns for each number of acetylated sites, analogous to the procedure we used for FUS LC phosphorylation above. We found that with increasing number of modifications, the chain dimension increases only slightly, from 3.2 nm to 3.3 nm (**Figure 2C, D**). Importantly, sequences carrying the same number of acetylated lysines may exhibit the same degree of variation of chain dimensions based on different arrangements of the acetylation sites. We note that in this case, the sequences with minimized (maximized) SCD achieve significantly lower (higher) *R*_g_ values as would be expected (**Figure 2D**).

Unlike FUS LC, the variability in the chain dimensions for a fixed number of modified sites is large as compared to the changes associated with increasing the number of modified sites. While FUS will generally become more expanded upon addition of PTMs, the single chain properties of DDX3 may change more drastically upon rearrangement of PTMs, than upon addition.

### Comparing post-translationally modified residues to common modification mimetics

A widely used approach in experimental studies of post-translational modifications is to mimic the change in amino acid character caused by the modification with a natural amino acid substitution. This allows for precise control of the location of modified residues. In the case of phosphorylation, serine and threonine are typically mutated to aspartic acid(56) or glutamic acid(48, 57, 58) which mimic the negative charge and larger size of the phosphate group. Here we evaluate aspartic acid or glutamic acid substitutions as phosphomimics using our model. We found that aspartic acid and glutamic acid do not fully recapitulate the chain dimensions observed with phosphorylation (**Figure 3A**), likely because phosphorylation results in a residue with −2 charge at neutral pH compared to the −1 charge of the phosphomimics. We tested this hypothesis that the dominant contribution to the difference in chain expansion for phosphorylation vs phosphomimetic mutation is due to the difference in added charge by replotting *R*_g_ as a function of net charge (**Figure 3B**). Indeed, we find that the effect of phosphorylation on FUS LC chain dimensions in the coarse-grained model appears to be largely driven by the charge of the modifications.

**Figure 3:**
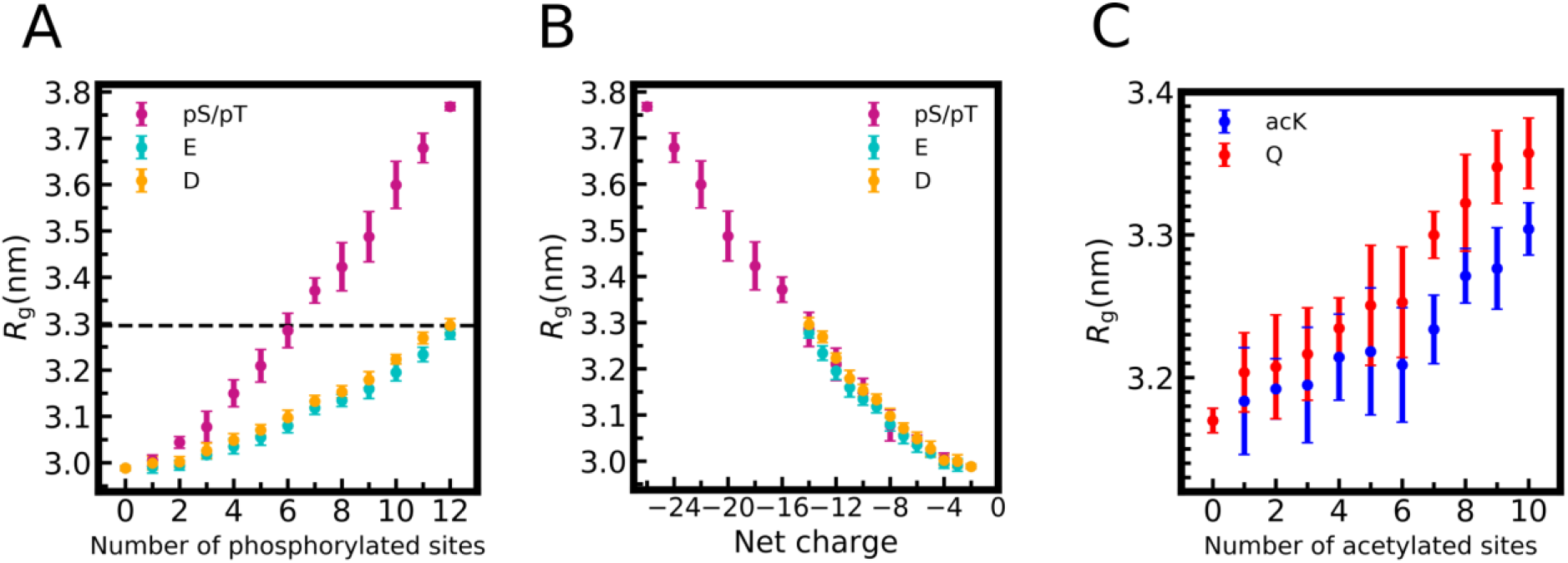
Comparing post-translationally modified residues to common modification mimetics. **A)** Effect of FUS LC phosphomimetic substitutions glutamic acid (E, cyan) or aspartic acid (D, orange) on *R*_g_ compared to phosphorylation (pS/pT,violet)**. B)** The same data as in **A)** but with x-axis shown as net charge of FUS LC, demonstrating that the difference between phosphomimetic and phosphorylated *R*_g_ can be largely explained by difference in net charge of aspartic acid and glutamine acid (−1) and phosphoserine or phosphothreonine (−2). **C)** Effect of DDX3X IDR acetylmimetic glutamine (Q, red) substitutions on *R*_g_ compared to acetylated lysine (acK, blue). Data represent mean and standard deviation from 10 individual simulations with random phosphorylation or substitution patterns per site except for 0 or n=max number of substitutions (for which there is only one pattern – all or none) where the block average and SEM is shown.

To mimic acetylation of lysine which removes the positive charge of the amino acid, researchers often mutate lysine to the uncharged, polar amino acid glutamine(59, 60). In our model, we find that glutamine substitutions mimic the changes in chain expansion caused by lysine acetylation in DDX3X IDR1 reasonably well (**Figure 3C**).

### Different PTM patterns with the same number of modifications have differing effects on LLPS

We next tested effect of the placement of PTMs on LLPS. We noted earlier that there may be dramatic differences in the single chain properties for sequences with the same number of PTMs. To understand the differences between the sequences, we decided to focus on DDX3X IDR1 where, even when the number of modifications is held constant, the arrangement of the acetylation modifications may have a large impact on *R*_g_ (**Figure 2D**). For the two sequences with highest or lowest values of *R*_g_ with a set number of modifications based on SCD, we computed *T_θ_* and found that *T_θ_* is significantly different for sequences with the same modification number (and hence same net charge) but different sequence (**Figure 4A**). Given that *T_θ_* (and *R*_g_) of an IDR is correlated with phase separation behavior(61), we sought to test how the placement of PTMs impacts the phase diagram for phase separation in the coarse-grained model. To demonstrate the position-specific effect of PTMs on the phase diagram of acetylated DDX3X IDR1, we simulated the two sequences with 4 acetylated lysines that exhibited the greatest difference in single chain properties. We find that positioning 4 acetylated lysines within the predominantly negatively charged C-terminus of DDX3X IDR1 enhances LLPS compared to the sequence that places the 4 acetylated lysines in the polyampholytic N-terminal/central region (**Figure 4B**). Together, these data demonstrate that the arrangement as well as the quantity of PTMs can have a large impact on the phase separation behavior of IDRs. This also highlights the necessity of methods that allow for control over which PTMs are “activated” in order to fully characterize a disordered protein and its range of responses to regulation by post-translational modifications.

**Figure 4.**
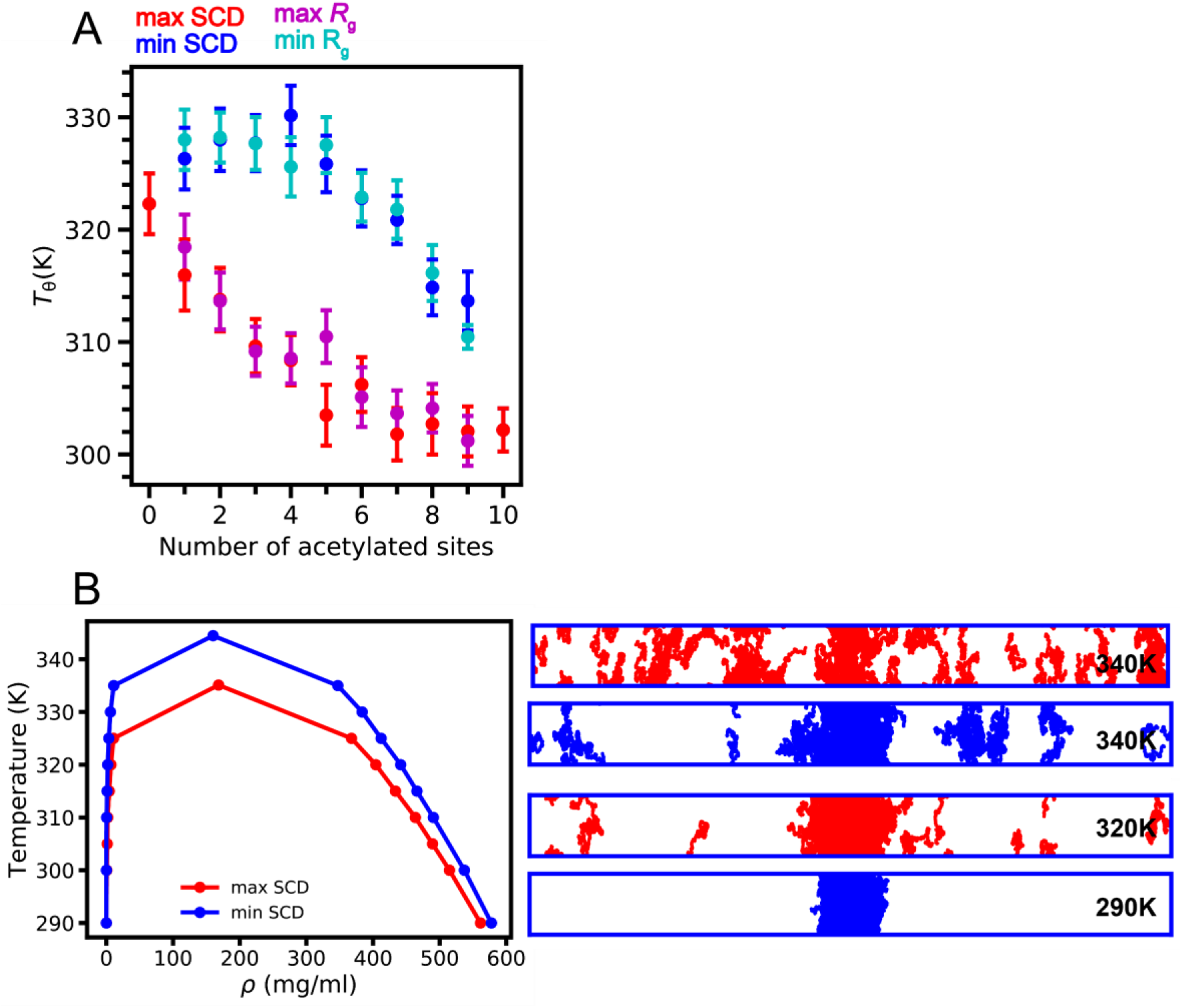
Different PTM patterns with the same number of modifications have differing effects on chain expansion and LLPS. **A)** *T_θ_* temperature of *R*_g_ minimum (cyan) and *R*_g_ maximum (magenta), SCD min (blue) and SCD max (red) DDX3X IDR1 sequences acetylated at n=1 to n=9 sites. Non-acetylated (0) and fully acetylated (10) variants are shown in green. **B)** Phase diagram of SCD min and SCD max of DDX3X IDR1 acetylated at 4 lysines are shown along with the simulation snapshots at multiple temperatures.

### Predicting effect of different modification patterns on phase separation: comparison to experiment

To test the predictive ability of the model, we decided to further explore the differences in predicted phase behavior for different modification patterns at a set number of modifications. To this end, we examined a series of single-position acetyl-mimetics (lysine to glutamine substitutions) of DDX3X IDR1 whose phase separation has been previously examined by experiment(48). We again show that in the coarse-grained model, single glutamine mimetics and acetylated lysine residues at the same position yield similar results (**Figure 5A**). However, between different single modifications we observe widely varying effects on chain properties, highlighting the importance of not only the number of PTMs, but the location (**Figure 5A**). Given that *R*_g_ is an excellent predictor of IDR phase separation(62) we then quantified the correlation between computationally predicted *R*_g_ and previously published data on phase separation measured experimentally by turbidity. As expected, none of these single position modifications has as much impact as simultaneous acetylmimetic substitutions at all 10 lysine positions (**Figure 5B**). Importantly, *R*_g_ is negatively correlated with the extent of phase separation as measured by experimental turbidity values (**Figure 5B**)(48). Thus, our findings demonstrate that this coarse-grained model provides predictive information regarding the effect of PTM location on the phase separation of IDRs.

**Figure 5:**
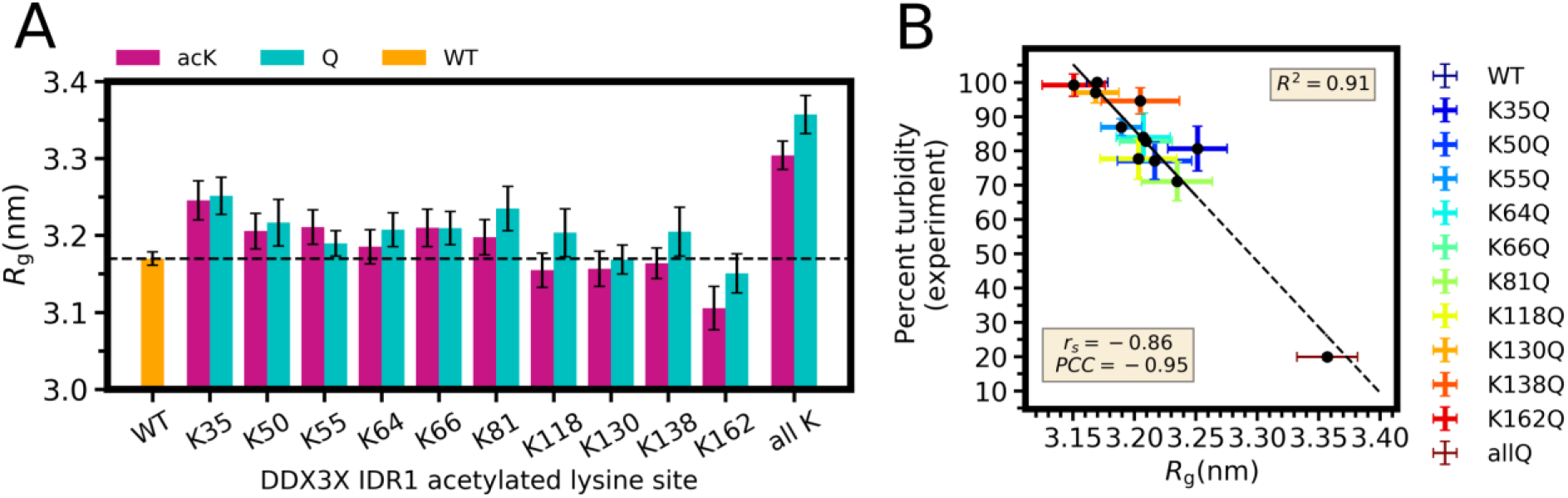
Predicting effect of different modification patterns on phase separation: comparison to experiment. **A)** Lysine acetylation (acK, violet) and lysine to glutamine acetyl mimetic mutants (cyan) show a range of *R*_g_ values compared to WT (orange) and all fully acetylated (all K) or fully acetyl mimetic (allQ). **B)** Correlation of percent turbidity measured by experiment and *R*_g_ (which is related to the phase separation behavior) predicted from the simulations. The values of spearman’s (rank order, r_s_) and Pearson’s coefficient (PCC) show strong negative monotonic and linear correlation, respectively.

## Discussion

Understanding the role of PTMs on phase separation is a critical element to controlling membraneless organelle formation in cells and engineering LLPS *in vitro*. In the present study we found that phosphorylation increases chain dimensions of FUS LC quickly and monotonically but that acetylation of DDX3X IDR1 has only a small average effect on chain expansion. Coarse-grained single-chain simulations revealed that phosphorylation expands FUS LC more drastically than phosphomimetic mutations because of the higher net charge of phosphorylation compared to the phosphomimetic substitutions. We found that particular patterns of PTMs with the same number of modified sites result in a large diversity in chain expansion which is strongly correlated with the propensity to phase separate. Importantly, we show modification patterns with the same number of modified sites can have dramatically different phase separation depending on the position of the modified residues. Furthermore, the model can predict the effect of position-specific modification on experimentally observed DDX3X IDR phase separation. Together our data suggest that the response of different disordered regions to multi-site modification depends on the modification patterning. In the future, it will be interesting to probe the effect on phase separation of IDR ensembles containing heterogenous combinations of IDRs with a varied distribution of both number and positions of modifications.

Our findings highlight the importance of combinatorial effects exerted by phosphorylation and acetylation on disordered proteins. Such effects typically remain unexplored by *in vitro* biochemistry approaches because protein modification reactions targeting only certain modification sites are difficult to achieve. However, we have noted that DNA-dependent protein kinase targets 12 SQ or TQ sites for phosphorylation with high specificity, modifying only these 12 out of a total of 52 S/T sites in FUS LC(25). With our coarse-grained model we can selectively modify amino acids of disordered regions and predict their effect on single-chain properties and phase behavior. These simulations can then enable future engineering of site-specific phosphorylation by introducing, for example, only SQ/TQ sites (and not other S/T sites) as specific targets for DNA-PK modification. Such computational studies can also aid predictions of phase separation properties of with IDRs with multi-site modifications.

We demonstrated that charge-changing PTMs lysine acetylation and serine/threonine/tyrosine phosphorylation can have a large and position-specific effect on IDR LLPS. It should be noted that threonine/serine phosphorylation and lysine acetylation are not the only PTMs with known or potential effect on LLPS. Methylation of arginine residues in proteins leads to stimulation(63) or suppression(8, 37, 64)of phase separation depending on the sequence content and the underlying cellular function. Importantly, methylation of arginine (and lysine) residues does not alter the charge of the residue (unlike lysine acetylation and serine/threonine/tyrosine phosphorylation). Hence, interactions other than charge are likely responsible for the changes to phase behavior by methylation and may be challenging to capture in coarse-grained model frameworks. Investigation of the role of methylated arginines and lysines on LLPS using transferable coarse-grained models remains important future work.

## Methods

### Hydropathy scale model for post translational modifications

The functional form of the total energy of the system is

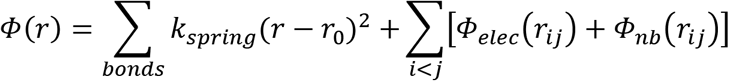

where *Φ_bond_* is a standard harmonic spring with 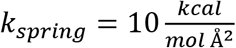 and *r*_0_ = 3.8 Å. The screened electrostatic term is represented using a Debye-Hückel type form:

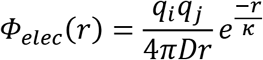

where *q_i_* and *q_j_* are the net charges of formally charged amino acids (D,E = −1; K,R = 1; H = 0.5), *D* is the dielectric constant, which is set to 80 for water, and κ is the screening length which is set to 10 Å to represent a salt concentration of about 100 mM. The non-bonded interactions are described by the Ashbaugh-Hatch functional form which has been previously applied to the study of disordered proteins(65). In this Lennard-Jones-like functional form, the attractiveness of the interactions is scaled by the arithmetic average of hydropathy for the two interacting amino acid types, *λ* = 0.5(*λ*_*i*_ + *λ*_*j*_).

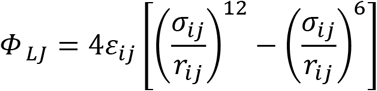

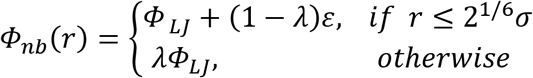

where *Φ_LJ_* is the standard Lennard-Jones potential and *σ* is the arithmetic average of the vdW radius:

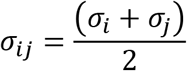

where *σ* = 2*r*_*vdw*_ since the atoms are modeled as spheres, and their impenetrable volumes are defined by the vdW radius(66) *r*_*vdw*_.

### Determining hydropathy values from atomic charges

The original HPS model is based on the hydrophobicity scale put forth by Kapcha and Rossky(49) using the atomic charges from the atomistic OPLS force field for proteins. The hydropathy scale was then scaled to the range of 0 to 1 with 0 being the least hydrophobic and 1 being the most hydrophobic. We further extend this procedure to obtain hydropathy values for post-translationally modified amino acid residues using partial atomic charges from the FF-PTM force field(50). Atoms with a partial charge magnitude ranging from 0 to 0.25 are considered non polar and are assigned a hydropathy value of −1. In the binary atomic-level hydropathy scale, atoms with a partial charge magnitude greater than 0.25 are considered polar and are assigned a hydropathy value of +1. The residue-level hydropathy values are calculated as a weighted sum of the atomic hydropathy values according to the following rules: hydrophobic (non-polar) atoms contribute −0.5, hydrophilic (polar) atoms contribute +1 and charged atoms contribute +2. An atom in charged residues is classified as charged if it belongs to the terminal polar group where the magnitude of the sum of all atomic partial charges in that group is greater than 0.5(49).

### Simulations framework

In order to obtain the temperature under which a protein chain behaves as an ideal solvent (*T_θ_*), single-chain simulations were conducted for 1 μs at a range of temperatures using replica exchange molecular dynamics (REMD)(51) with a temperature list of 150.0, 170.1, 193.0, 218.9, 248.3, 281.7, 300, 362.4, 411.1, 466.3, 529.0, and 600.0 K to enhance ergodic sampling of the IDR conformational ensemble(67). Simulations were conducted in cubic boxes with periodic boundaries, large enough that a protein chain will not encounter its periodic image, and temperature was maintained using a Langevin thermostat. All single-chain simulations were conducted using LAMMPS(68).

Coexistence simulations were conducted using slab geometry(69, 70) as in our previous work(16, 37, 44, 61, 71, 72). Simulations were conducted on 100 chains of FUS or DDX3 using the HPS model and the new parameters for PTMs. Simulations were conducted using HOOMD-Blue v2.1.5(73) for 5 μs each in serial with a Langevin thermostat, where the first μs was discarded as equilibration.

### Single molecule properties

Using the radius of gyration (*R*_g_) and the polymer scaling exponent (*ν*) in different temperatures as descriptors of the phase behavior, we calculate the temperature at which a protein behaves as an ideal polymer; the point where protein-protein interactions and protein-solvent interactions become energetically equally favorable (*T_θ_*)(52). For each temperature we estimated the polymer scaling exponent (*ν*) by fitting to:

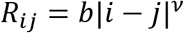

where *b* is the Kuhn length, which is set to 0.55 nm for disordered proteins(74) and *R_ij_* the intrachain separation defined as the distance between residue i and residue j. Next, we interpolated the results to find the temperature at which v = 0.5 which is correlated with the critical temperature for phase separation(61).

### Phase diagrams

Phase diagrams were extracted from coexistence simulations by measuring the density of protein in the dense phase in the center, and in the low-density phase outside. The critical temperature and critical density were obtained using the top 7 highest temperatures for the max SCD case and 8 for the min SCD case in which we observed phase coexistence. We fit the coexistence densities at these temperatures to a fitting function:

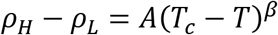

where β is set to 0.325 according to the universality class of the 3D Ising model(73).

## Author contributions

T.M.P and J.M designed and performed single-chain coarse-grained simulations and analyzed the resulting data. N.J and J.M designed, performed and analyzed slab simulations. G.L.D, Y.C.K and J.M developed the methodology, provided the software and supervised the formal analysis. J.M and N.L.F contributed to project administration and funding acquisition. T.M.P, N.L.F and J.M wrote the manuscript with comments from all authors.

## Acknowledgements

Research was supported in part by NIGMS R01GM118530 (to N.L.F.), NINDS and NIA R01NS116176 (to N.L.F. and J.M.), NSF 2004796 (to J.M.), and NSF 1845734 (to N.L.F.). Y.C.K. is supported by the Office of Naval Research via the U.S. Naval Research Laboratory base program. Use of the high-performance computing capabilities of the Extreme Science and Engineering Discovery Environment (XSEDE), which is supported by the NSF grant TG-MCB-120014, is gratefully acknowledged. The content is solely the responsibility of the authors and does not necessarily represent the official views of the funding agencies. We thank Anastasia Murthy and Veronica Ryan for helpful discussions. We thank Patrick Matthias for providing previously published tabulated data on DDX3X phase separation.

## References

1. Shin, Y., and C.P. Brangwynne. 2017. Liquid phase condensation in cell physiology and disease. Science (80-.). 357.

2. Ditlev, J.A., L.B. Case, and M.K. Rosen. 2018. Who’s In and Who’s Out— Compositional Control of Biomolecular Condensates. J. Mol. Biol. 430:4666–4684.

3. Ryan, V.H., and N.L. Fawzi. 2019. Physiological, Pathological, and Targetable Membraneless Organelles in Neurons. Trends Neurosci. 42:693–708.

4. Banani, S.F., H.O. Lee, A.A. Hyman, and M.K. Rosen. 2017. Biomolecular condensates: Organizers of cellular biochemistry. Nat. Rev. Mol. Cell Biol. 18.

5. Molliex, A., J. Temirov, J. Lee, M. Coughlin, A.P. Kanagaraj, H.J. Kim, T. Mittag, and J.P. Taylor. 2015. Phase Separation by Low Complexity Domains Promotes Stress Granule Assembly and Drives Pathological Fibrillization. Cell. 163:123–133.

6. Feric, M., N. Vaidya, T.S. Harmon, R.W. Kriwacki, R. V Pappu, C.P.B. Correspondence, D.M. Mitrea, L. Zhu, T.M. Richardson, and C.P. Brangwynne. 2016. Coexisting Liquid Phases Underlie Nucleolar Subcompartments Article Coexisting Liquid Phases Underlie Nucleolar Subcompartments. Cell. 165:1686–1697.

7. Darling, A.L., Y. Liu, C.J. Oldfield, and V.N. Uversky. 2018. Intrinsically Disordered Proteome of Human Membrane-Less Organelles. Proteomics. 18:1–12.

8. Qamar, S., G.Z. Wang, S.J. Randle, F.S. Ruggeri, J.A. Varela, J.Q. Lin, E.C. Phillips, A. Miyashita, D. Williams, F. Ströhl, W. Meadows, R. Ferry, V.J. Dardov, G.G. Tartaglia, L.A. Farrer, G.S. Kaminski Schierle, C.F. Kaminski, C.E. Holt, P.E. Fraser, G. Schmitt-Ulms, D. Klenerman, T. Knowles, M. Vendruscolo, and P. St George-Hyslop. 2018. FUS Phase Separation Is Modulated by a Molecular Chaperone and Methylation of Arginine Cation-π Interactions. Cell. 173:720–734.e15.

9. Vernon, R.M., P.A. Chong, B. Tsang, T.H. Kim, A. Bah, P. Farber, H. Lin, and J.D. Forman-Kay. 2018. Pi-Pi contacts are an overlooked protein feature relevant to phase separation. Elife. 7:1–48.

10. Burke, K.A., A.M. Janke, C.L. Rhine, and N.L. Fawzi. 2015. Residue-by-Residue View of In Vitro FUS Granules that Bind the C-Terminal Domain of RNA Polymerase II. Mol. Cell. 60:231–241.

11. Dignon, G.L., R.B. Best, and J. Mittal. 2020. Biomolecular Phase Separation: From Molecular Driving Forces to Macroscopic Properties. Annu. Rev. Phys. Chem. 71:1–23.

12. Murthy, A.C., G.L. Dignon, Y. Kan, G.H. Zerze, S.H. Parekh, J. Mittal, and N.L. Fawzi. 2019. Molecular interactions underlying liquid–liquid phase separation of the FUS low-complexity domain. Nat. Struct. Mol. Biol. 26:637–648.

13. Schwartz, J.C., X. Wang, E.R. Podell, and T.R. Cech. 2013. RNA Seeds Higher-Order Assembly of FUS Protein. Cell Rep. 5:918–925.

14. Lin, Y., J. McCarty, J.N. Rauch, K.T. Delaney, K.S. Kosik, G.H. Fredrickson, J.E. Shea, and S. Han. 2019. Narrow equilibrium window for complex coacervation of tau and RNA under cellular conditions. Elife. 8:1–31.

15. Hayes, M.H., E.H. Peuchen, N.J. Dovichi, and D.L. Weeks. 2018. Dual roles for ATP in the regulation of phase separated protein aggregates in xenopus oocyte nucleoli. Elife. 7:1–24.

16. Dignon, G.L., W. Zheng, Y.C. Kim, and J. Mittal. 2019. Temperature-Controlled Liquid-Liquid Phase Separation of Disordered Proteins. ACS Cent. Sci. 0002:821–830.

17. Hofweber, M., and D. Dormann. 2019. Friend or foe-Post-translational modifications as regulators of phase separation and RNP granule dynamics. J. Biol. Chem. 294:7137–7150.

18. Bah, A., and J.D. Forman-Kay. 2016. Modulation of intrinsically disordered protein function by post-translational modifications. J. Biol. Chem. 291:6696–6705.

19. Khoury, G.A., R.C. Baliban, and C.A. Floudas. 2014. Proteome-wide post-translational modification statistics: Frequency analysis and curation of the swiss-prot database. Sci. Rep. 1:1–5.

20. Andrusiak, M.G., P. Sharifnia, X. Lyu, Z. Wang, A.M. Dickey, Z. Wu, A.D. Chisholm, and Y. Jin. 2019. Inhibition of Axon Regeneration by Liquid-like TIAR-2 Granules. Neuron. 104:290–304.e8.

21. Ferreon, J., A. Jain, K.-J. Choi, P. Tsoi, K. MacKenzie, S. Jung, and A. Ferreon. 2018. Acetylation Disfavors Tau Phase Separation. Int. J. Mol. Sci. 19:1360.

22. Audagnotto, M., and M. Dal Peraro. 2017. Protein post-translational modifications: In silico prediction tools and molecular modeling. Comput. Struct. Biotechnol. J. 15:307–319.

23. Kim, T.H., B. Tsang, R.M. Vernon, N. Sonenberg, L.E. Kay, and J.D. Forman-Kay. 2019. Phospho-dependent phase separation of FMRP and CAPRIN1 recapitulates regulation of translation and deadenylation. Science (80-.). 365:825–829.

24. Brunk, E., R.L. Chang, J. Xia, H. Hefzi, J.T. Yurkovich, D. Kim, E. Buckmiller, H.H. Wang, B.-K. Cho, C. Yang, B.O. Palsson, G.M. Church, N.E. Lewis, E. Brunk, N.E.L. Designed,; E Brunk, E. Buckmiller, and N.E.L. Performed. 2018. Characterizing posttranslational modifications in prokaryotic metabolism using a multiscale workflow. 115:11096–11101.

25. Monahan, Z., V.H. Ryan, A.M. Janke, K.A. Burke, S.N. Rhoads, G.H. Zerze, R. O’Meally, G.L. Dignon, A.E. Conicella, W. Zheng, R.B. Best, R.N. Cole, J. Mittal, F. Shewmaker, and N.L. Fawzi. 2017. Phosphorylation of the FUS low complexity domain disrupts phase separation, aggregation, and toxicity. EMBO J. 36:e201696394.

26. Tsang, B., J. Arsenault, R.M. Vernon, H. Lin, N. Sonenberg, L.-Y. Wang, A. Bah, and J.D. Forman-Kay. 2019. Phosphoregulated FMRP phase separation models activity-dependent translation through bidirectional control of mRNA granule formation. Proc. Natl. Acad. Sci. 201814385.

27. Snead, W.T., and A.S. Gladfelter. 2019. The Control Centers of Biomolecular Phase Separation: How Membrane Surfaces, PTMs, and Active Processes Regulate Condensation. Mol. Cell. 76:295–305.

28. Kamah, A., I. Huvent, F.X. Cantrelle, H. Qi, G. Lippens, I. Landrieu, and C. Smet-Nocca. 2014. Nuclear magnetic resonance analysis of the acetylation pattern of the neuronal Tau protein. Biochemistry. 53:3020–3032.

29. Wegmann, S., B. Eftekharzadeh, K. Tepper, K.M. Zoltowska, R.E. Bennett, S. Dujardin, P.R. Laskowski, D. MacKenzie, T. Kamath, C. Commins, C. Vanderburg, A.D. Roe, Z. Fan, A.M. Molliex, A. Hernandez Vega, D. Muller, A.A. Hyman, E. Mandelkow, J.P. Taylor, and B.T. Hyman. 2018. Tau protein liquid–liquid phase separation can initiate tau aggregation. EMBO J. e98049.

30. Martin, E.W., A.S. Holehouse, C.R. Grace, A. Hughes, R.V. Pappu, and T. Mittag. 2016. Sequence Determinants of the Conformational Properties of an Intrinsically Disordered Protein Prior to and upon Multisite Phosphorylation. J. Am. Chem. Soc. 138:15323–15335.

31. Huihui, J., T. Firman, and K. Ghosh. 2018. Modulating charge patterning and ionic strength as a strategy to induce conformational changes in intrinsically disordered proteins. J. Chem. Phys. 149.

32. Margreitter, C., D. Petrov, and B. Zagrovic. 2013. Vienna-PTM web server: a toolkit for MD simulations of protein post-translational modifications. Nucleic Acids Res. 41:422–426.

33. Rani, L., J. Mittal, and S.S. Mallajosyula. 2020. Effect of Phosphorylation and O-GlcNAcylation on Proline-Rich Domains of Tau. J. Phys. Chem. B. 124:1909–1918.

34. Khoury, G.A., J. Smadbeck, P. Tamamis, A.C. Vandris, C.A. Kieslich, and C.A. Floudas. 2014. Forcefield-NCAA: Ab initio charge parameters to aid in the discovery and design of therapeutic proteins and peptides with unnatural amino acids and their application to complement inhibitors of the compstatin family. ACS Synth. Biol. 3:855–869.

35. Khoury, G.A., J.P. Thompson, J. Smadbeck, C.A. Kieslich, and C.A. Floudas. 2013. Forcefield-PTM: Ab initio charge and AMBER forcefield parameters for frequently occurring post-translational modifications. J. Chem. Theory Comput. 9:5653–5674.

36. Zerze, G.H., and J. Mittal. 2015. Effect of O-Linked Glycosylation on the Equilibrium Structural Ensemble of Intrinsically Disordered Polypeptides. J. Phys. Chem. B. 119:15583–15592.

37. Ryan, V.H., G.L. Dignon, G.H. Zerze, C.V. Chabata, R. Silva, A.E. Conicella, J. Amaya, K.A. Burke, J. Mittal, and N.L. Fawzi. 2018. Mechanistic View of hnRNPA2 Low-Complexity Domain Structure, Interactions, and Phase Separation Altered by Mutation and Arginine Methylation. Mol. Cell. 69:465–479.e7.

38. Ryan, V.H., T.M. Perdikari, M.T. Naik, C.F. Saueressig, J. Lins, G.L. Dignon, J. Mittal, A.C. Hart, and N.L. Fawzi. 2020. Tyrosine phosphorylation regulates hnRNPA2 granule protein partitioning & reduces neurodegeneration. bioarxiv. 1–38.

39. Vymětal, J., V. Jurásková, and J. Vondrášek. 2019. AMBER and CHARMM Force Fields Inconsistently Portray the Microscopic Details of Phosphorylation. J. Chem. Theory Comput. 15:665–679.

40. Zheng, W., R.B. Best, J. Mittal, G.H. Zerze, W. Zheng, R.B. Best, and J. Mittal. 2019. Evolution of All-Atom Protein Force Fields to Improve Local and Global Properties. J. Phys. Chem. Lett. 10:2227–2234.

41. Saunders, M.G., and G.A. Voth. 2013. Coarse-Graining Methods for Computational Biology. Annu. Rev. Biophys. 42:73–93.

42. Ruff, K.M., R.V. Pappu, and A.S. Holehouse. 2019. Conformational preferences and phase behavior of intrinsically disordered low complexity sequences: insights from multiscale simulations. Curr. Opin. Struct. Biol. 56:1–10.

43. Dignon, G.L., W. Zheng, and J. Mittal. 2019. Simulation methods for liquid–liquid phase separation of disordered proteins. Curr. Opin. Chem. Eng. 23:92–98.

44. Dignon, G.L., W. Zheng, Y.C. Kim, R.B. Best, and J. Mittal. 2018. Sequence determinants of protein phase behavior from a coarse-grained model. PLoS Comput. Biol. 14.

45. Sieradzan, A.K., M. Bogunia, P. Mech, R. Ganzynkowicz, A. Giełdoń, A. Liwo, and M. Makowski. 2019. Introduction of Phosphorylated Residues into the UNRES Coarse-Grained Model: Toward Modeling of Signaling Processes. J. Phys. Chem. B. 123:5721–5729.

46. Shen, T., C. Zong, D. Hamelberg, J. Andrew Mccammon, and P.G. Wolynes. 2005. The folding energy landscape and phosphorylation: modeling the conformational switch of the NFAT regulatory domain. FASEB J. 19:1389–1395.

47. Rhoads, S.N., Z.T. Monahan, D.S. Yee, and F.P. Shewmaker. 2018. The Role of Post-Translational Modifications on Prion-Like Aggregation and Liquid-Phase Separation of FUS. Int. J. Mol. Sci. 19.

48. Saito, M., D. Hess, J. Eglinger, A.W. Fritsch, M. Kreysing, B.T. Weinert, C. Choudhary, and P. Matthias. 2019. Acetylation of intrinsically disordered regions regulates phase separation. Nat. Chem. Bio. 15:51–61.

49. Kapcha, L.H., and P.J. Rossky. 2014. A simple atomic-level hydrophobicity scale reveals protein interfacial structure. J. Mol. Biol. 426:484–498.

50. Khoury, G.A., J.P. Thompson, J. Smadbeck, C.A. Kieslich, and C.A. Floudas. 2013. Forcefield_PTM: Ab Initio Charge and AMBER Forcefield Parameters for Frequently Occurring Post-Translational Modifications. J. Chem. Theory Comput. 9:5653–5674.

51. Sugita Y., O.Y. Y. Sugita, and Y. Okamoto. 1999. Replica-exchange molecular dynamics method for protein folding. Chem. Phys. Lett. 314:141.

52. Flory, P.J. 1949. Statistical Mechanics of Cross-Linked Polymer Networks I. Stat. Mech. Dilute Polym. Solut. J. Chem. Phys. 17:1347.

53. Firman, T., and K. Ghosh. 2018. Sequence charge decoration dictates coil-globule transition in intrinsically disordered proteins. J. Chem. Phys. 148.

54. Zheng, W., G.L. Dignon, M. Brown, Y.C. Kim, and J. Mittal. 2020. Hydropathy Patterning Complements Charge Patterning to Describe Conformational Preferences of Disordered Proteins. J. Phys. Chem. Lett.

55. Das, S., A.N. Amin, Y.H. Lin, and H.S. Chan. 2018. Coarse-grained residue-based models of disordered protein condensates: utility and limitations of simple charge pattern parameters. Phys. Chem. Chem. Phys. 20:28558–28574.

56. Brady, O.A., P. Meng, Y. Zheng, Y. Mao, and F. Hu. 2011. Regulation of TDP-43 aggregation by phosphorylation andp62/SQSTM1. J. Neurochem. 116:248–259.

57. Dephoure, N., K.L. Gould, S.P. Gygi, and D.R. Kellogg. 2013. Mapping and analysis of phosphorylation sites: A quick guide for cell biologists. Mol. Biol. Cell. 24:535–542.

58. Oulhen, N., S. Boulben, M. Bidinosti, J. Morales, P. Cormier, and B. Cosson. 2009. A variant mimicking hyperphosphorylated 4E-BP inhibits protein synthesis in a sea urchin cell-free, cap-dependent translation system. PLoS One. 4:e5070.

59. Li, M., J. Luo, C.L. Brooks, and W. Gu. 2002. Acetylation of p53 inhibits its ubiquitination by Mdm2. J. Biol. Chem. 277:50607–50611.

60. Alaei, S.R., C.K. Abrams, J.C. Bulinski, E.L. Hertzberg, and M.M. Freidin. 2018. Acetylation of C-terminal lysines modulates protein turnover and stability of Connexin-32. BMC Cell Biol. 19.

61. Dignon, G.L., W. Zheng, R.B. Best, Y.C. Kim, and J. Mittal. 2018. Relation between single-molecule properties and phase behavior of intrinsically disordered proteins. Proc. Natl. Acad. Sci. 115:9929–9934.

62. Lin, Y.H., J.D. Forman-Kay, and H.S. Chan. 2018. Theories for Sequence-Dependent Phase Behaviors of Biomolecular Condensates. Biochemistry. 57:2499–2508.

63. Arribas-Layton, M., J. Dennis, E.J. Bennett, C.K. Damgaard, and J. Lykke-Andersen. 2016. The C-Terminal RGG Domain of Human Lsm4 Promotes Processing Body Formation Stimulated by Arginine Dimethylation. Mol. Cell. Biol. 36:2226–2235.

64. Matsumoto, K., H. Nakayama, M. Yoshimura, A. Masuda, N. Dohmae, S. Matsumoto, and M. Tsujimoto. 2012. PRMT1 is required for RAP55 to localize to processing bodies. RNA Biol. 9:610–623.

65. Ashbaugh, H.S., and H.W. Hatch. 2008. Natively unfolded protein stability as a coil-to-globule transition in charge/hydropathy space. J. Am. Chem. Soc. 130:9536–9542.

66. Creighton, T.E. Proteins: structures and molecular properties. Helevetian Press.

67. Hansmann, U.H.E. 1997. Parallel tempering algorithm for conformational studies of biological molecules. Chem. Phys. Lett. 281:140–150.

68. Plimpton, S. 1995. Fast Parallel Algorithms for Short-Range Molecular Dynamics. J. Comput. Phys. 117:1–19.

69. Blas, F.J., L.G. MacDowell, E. De Miguel, and G. Jackson. 2008. Vapor-liquid interfacial properties of fully flexible Lennard-Jones chains. J. Chem. Phys. 129.

70. Silmore, K.S., M.P. Howard, and A.Z. Panagiotopoulos. 2017. Vapour-liquid phase equilibrium and surface tension of fully flexible Lennard-Jones chains. Mol. Phys. 115:320–327.

71. Schuster, B.S., E.H. Reed, R. Parthasarathy, C.N. Jahnke, R.M. Caldwell, J.G. Bermudez, H. Ramage, M.C. Good, and D.A. Hammer. 2018. Controllable protein phase separation and modular recruitment to form responsive membraneless organelles. Nat. Commun. 9:1–12.

72. Schuster, B.S., G.L. Dignon, W.S. Tang, F.M. Kelley, A.K. Ranganath, C.N. Jahnke, A.G. Simpkins, R.M. Regy, D.A. Hammer, M.C. Good, and J. Mittal. 2020. Identifying sequence perturbations to an intrinsically disordered protein that determine its phase-separation behavior. Proc. Natl. Acad. Sci.

73. Anderson, J.A., C.D. Lorenz, and A. Travesset. 2008. General purpose molecular dynamics simulations fully implemented on graphics processing units. J. Comput. Phys. 227:5342–5359.

74. Hofmann, H., A. Soranno, A. Borgia, K. Gast, D. Nettels, and B. Schuler. 2012. Polymer scaling laws of unfolded and intrinsically disordered proteins quantified with single-molecule spectroscopy. Proc. Natl. Acad. Sci. 109:16155–16160.

